# Butterfly wings exhibit spatial variation in chromatin accessibility

**DOI:** 10.1101/2022.01.21.477190

**Authors:** Heidi Connahs, Mainak das Gupta, Antónia Monteiro

**Affiliations:** Department of Biological Sciences, National University of Singapore

**Keywords:** FAIRE-seq, Chromatin accessibility, Butterfly wings, Morphology, GC content, Hox genes, cofactors

## Abstract

Butterfly wings exhibit a diversity of patterns which can vary between forewings and hindwings and spatially across the same wing. Regulation of morphological variation involves changes in how genes are expressed across different spatial scales which is driven by chromatin dynamics during development. How patterns of chromatin dynamics correspond to morphological variation remains unclear. Here we compared the chromatin landscape between forewings and hindwings and also across the proximal and distal regions of the hindwings in two butterfly species, *Bicyclus anynana* and *Danaus plexippus*. We found that the chromatin profile varied significantly between the different wing regions, however, there was no clear correspondence between the chromatin profile and the wing patterns. In some cases, wing regions with different phenotypes shared the same chromatin profile whereas those with a similar phenotype had a different profile. We also found that in the forewing, open chromatin regions (OCRs) were AT rich whereas those in the hindwing were GC rich. GC content of the OCRs also varied between the proximal and hindwing regions. These differences in GC content were also reflected in the transcription factor binding motifs that were differentially enriched between the wings and wing regions. Our results suggest that distinct wing patterns may result from the interaction of pioneer factors, including Hox genes, differentially opening chromatin in different wings and wing regions and cooperating with other transcriptions factors, that show preferences for specific GC content, to function either as activator or repressors of nearby genes.

## Introduction

Butterflies are renowned for their striking wing patterns which can vary dramatically between forewings and hindwings and across different regions and surfaces of the wing. Numerous studies have characterized spatial patterns of gene expression across different regions of the wing (Ferguson and Jiggins 2009; Ferguson et al. 2011; Prakash and Monteiro 2018; Hanly et al. 2019; Banerjee and Monteiro 2020) and also identified genes that regulate specific colors and pattern elements (Martin and Reed, 2010; Reed et al., 2011; Kunte et al., 2014; Mazo-vargas et al., 2017; Matsuoka and Monteiro, 2018; Connahs et al., 2019). The spatial and temporal expression pattern of any gene is defined by chromatin dynamics in which specific regions of chromatin start to become open during development, via the binding of pioneer factors, facilitating the binding of further transcription factors to open chromatin regions (OCRs), which ultimately drive patterns of gene expression (Shlyueva et al. 2014). But precisely how these changes in chromatin dynamics translate to morphological variation is still poorly understood.

Most studies to date, which have examined the chromatin landscape of whole tissues, generally show that at the genome-wide level, chromatin accessibility is surprisingly invariant. Studies in the butterflies *Junonia* and *Heliconius*, have shown similar chromatin profiles between forewings and hindwings despite them displaying different color patterns (Lewis and Reed 2018; Burg et al. 2019; Lewis et al. 2019). In *Drosophila*, a comparison of the chromatin landscape across different appendages also revealed that the same enhancers were accessible at most genomic regions with the exception of the loci around master regulators (McKay and Lieb, 2014). Similar to the studies in *Junonia* and *Heliconius*, changes in the chromatin profile corresponded with different stages of development, whereas at the same developmental stage even tissues with very different phenotypes had a remarkably similar profile.

Based on these findings how can we explain the development of different morphologies if the chromatin landscape is largely identical? McKay and Leib suggested that master regulators, which were the few genes they found to be differentially accessible as well as expressed between different appendages in flies (e.g. *Ultrabithorax* (*Ubx*) and *vestigial*), are responsible for the morphological differences. These genes would bind to the same set of accessible enhancers of downstream genes effecting changes in gene expression across appendages. Differential chromatin accessibility for two key regulators was also found in *Junonia* and *Heliconius* butterflies (Lewis and Reed 2018; Burg et al. 2019; Lewis et al. 2019), where the only difference observed in the chromatin profile of forewings and hindwings was around the loci of *Ubx*, which defines hindwing identity (Weatherbee et al. 1999; Tomoyasu 2017; Matsuoka and Monteiro 2021) and *Abd-a* (*Heliconius* only) a neighboring gene to *Ubx* (Deutsch 2005). Other key genes such as *optix*, previously shown to regulate red wing pattern elements in *Heliconius*, did not have distinct chromatin profiles surrounding its OCRs between forewings and hindwings even when only one wing had a red pattern element (Lewis. et al. 2019). A ChIP-seq also showed that Optix protein itself binds the same OCRs in both wings, and deletion of OCRs containing *optix* binding sites leads to disruption of these patterns (Lewis et al. 2019). How information processed at those OCRs contribute to the differential expression of *optix* in forewings and hindwings remains, thus, unclear.

An alternative explanation to McKay and Leib’s suggestion is that different wing phenotypes do in fact correspond to changes in the chromatin profile but that capturing the chromatin profile at the whole tissue level is insufficient for detecting spatial differences occurring at a finer resolution. A study examining the chromatin profile across different regions of a *Drosophila* embryo showed distinct spatial differences in the chromatin landscape when compared to the profile obtained using whole embryos (Bozek et al. 2019). Examining the whole embryo resulted in an averaging out of the chromatin signal thus masking spatial differences in chromatin accessibility. These findings indicate that previous studies, using whole butterfly wings, or whole fly appendages, may have missed spatial differences in the chromatin landscape that could be associated with differences in wing/appendage phenotypes.

Here we examined whether the wing chromatin landscape corresponds to differences in wing color patterns. In the African butterfly, *Bicyclus anynana*, both forewings and hindwings display eyespots on the distal margin of the wings whereas the proximal wing region is largely homogeneous with few distinct pattern elements. We hypothesized that the chromatin profile would differ between the distal and proximal wing tissue, but would be more similar between the distal regions of the forewing and hindwing which have eyespots. We also compared the chromatin profile of the proximal and distal regions of the Monarch butterfly, *Danaus plexippus* which exhibits monochromatic spots along the distal margins of both wings and different patterning in more proximal wing regions. We then investigated whether these two butterflies shared regions of open chromatin that were highly conserved at the sequence level, and whether these regions exhibited functional conservation, i.e., were open in the same wings and/or wing regions compared to non-conserved OCRs. Finally, we performed a motif discovery analysis to examine whether the regions of open chromatin in different wings or wing regions were enriched in particular transcription factor binding motifs. This analysis aimed to discover transcription factors important in regulating differences in wing color pattern phenotypes.

## Results

### Open chromatin profile varies between different wings and wing regions

We first annotated the FAIRE-seq data using peaks observed in IGV, highlighting the significant peaks called from MACS2 to check whether open chromatin regions (OCRs) were open across all wing regions regardless of phenotype as observed in *Heliconius* and *Junonia*. For this analysis we focused on the significant peaks around 6 genes known to be involved in wing patterning (*Distal-less, Ecdysone receptor (EcR), spalt, decapentaplegic (dpp), wingless* and *engrailed*). For *B. anynana*, we compared the open chromatin landscape between the distal regions of the forewing (FWD) and hindwing (HWD) and also between the proximal and distal regions of the hindwings (HWP and HWD). For *D. plexippus* we compared the proximal and distal regions of the hindwings. In contrast to *B. anynana*, the data for the forewing (FW) represents the entire wing rather than just the distal region (Fig 1). For both butterfly species, we found that the chromatin landscape varied significantly between forewings and hindwings and between the proximal and distal regions of the hindwing (Figs 2–3, S1). We next examined the number of shared OCRs between the different wings and wing regions. For this analysis we used the FAIRE-seq peaks extracted from all scaffolds. For *D. plexippus*, 41,608 significant peaks were observed across the whole genome (FW = 21178, HWD=8911, HWP=11519). For *B. anynana*, we had data for 31 scaffolds and observed 629 significant peaks (FWD = 240, HWD=272, HWP=117). We found that in both species the majority of overlapping OCRs were observed between the HWP and FWD/FW regions which have very different wing phenotypes (Fig 4). In *D. plexippus* the fewest overlaps were observed between the FW and HWD regions. In *B. anynana*, we did not observe any peaks that were uniquely shared between the FWD and HWD regions despite the presence of eyespots in both of these wing regions (Fig 4).

**Fig 1.**
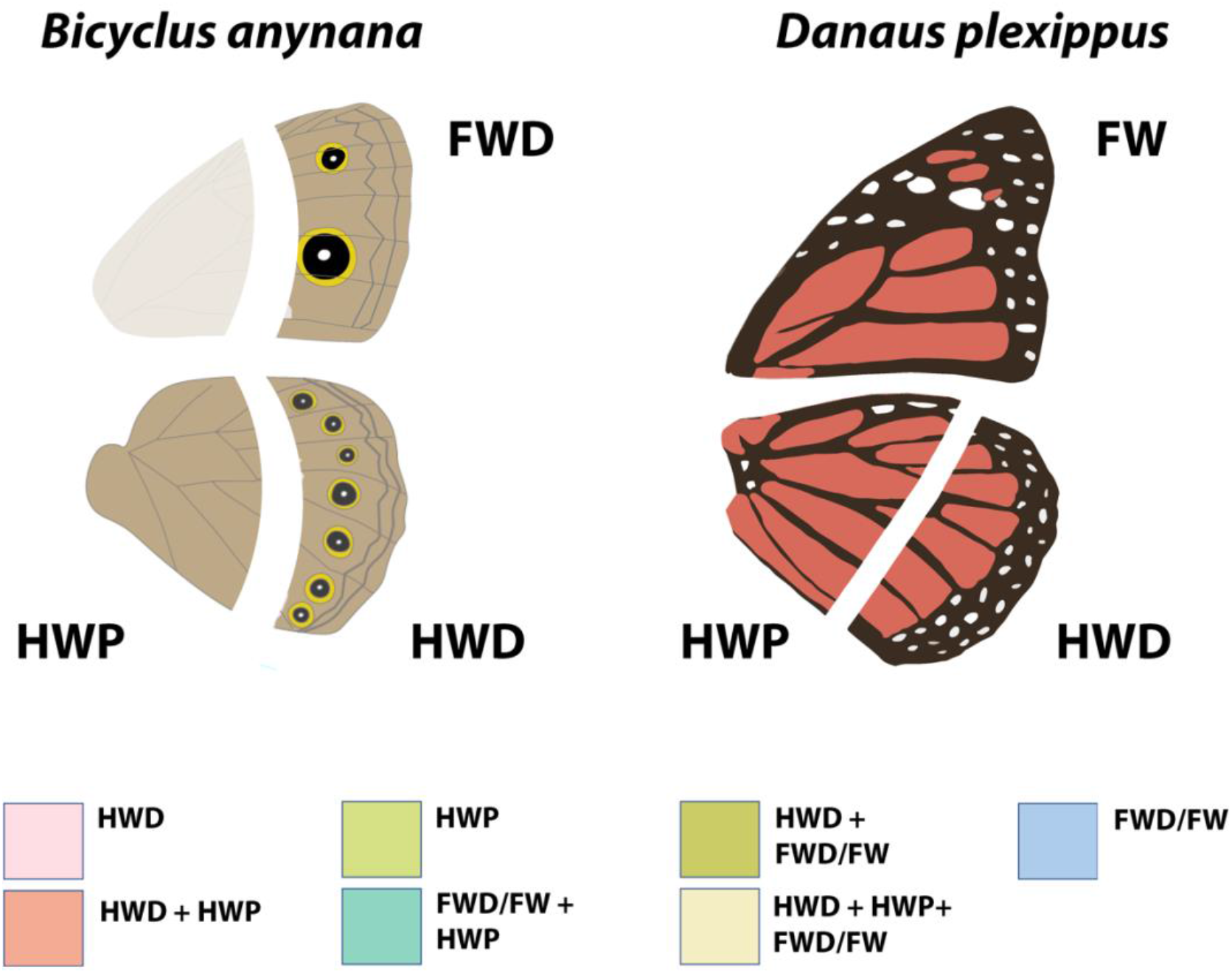
Wing dissections for FAIRE-seq. For *B. anynana*, libraries were prepared from the distal forewing (FWD), distal hindwing (HWD) and proximal hindwing (HWP). For *D. plexippus*, libraries were prepared for the entire forewing (FW), distal hindwing (HWD) and proximal hindwing (HWP). The color key below shows the color scheme that was used to code the significant peaks and those which are overlapping for the FAIRE-seq data (Figs 2–3, S1).

**Fig 2.**
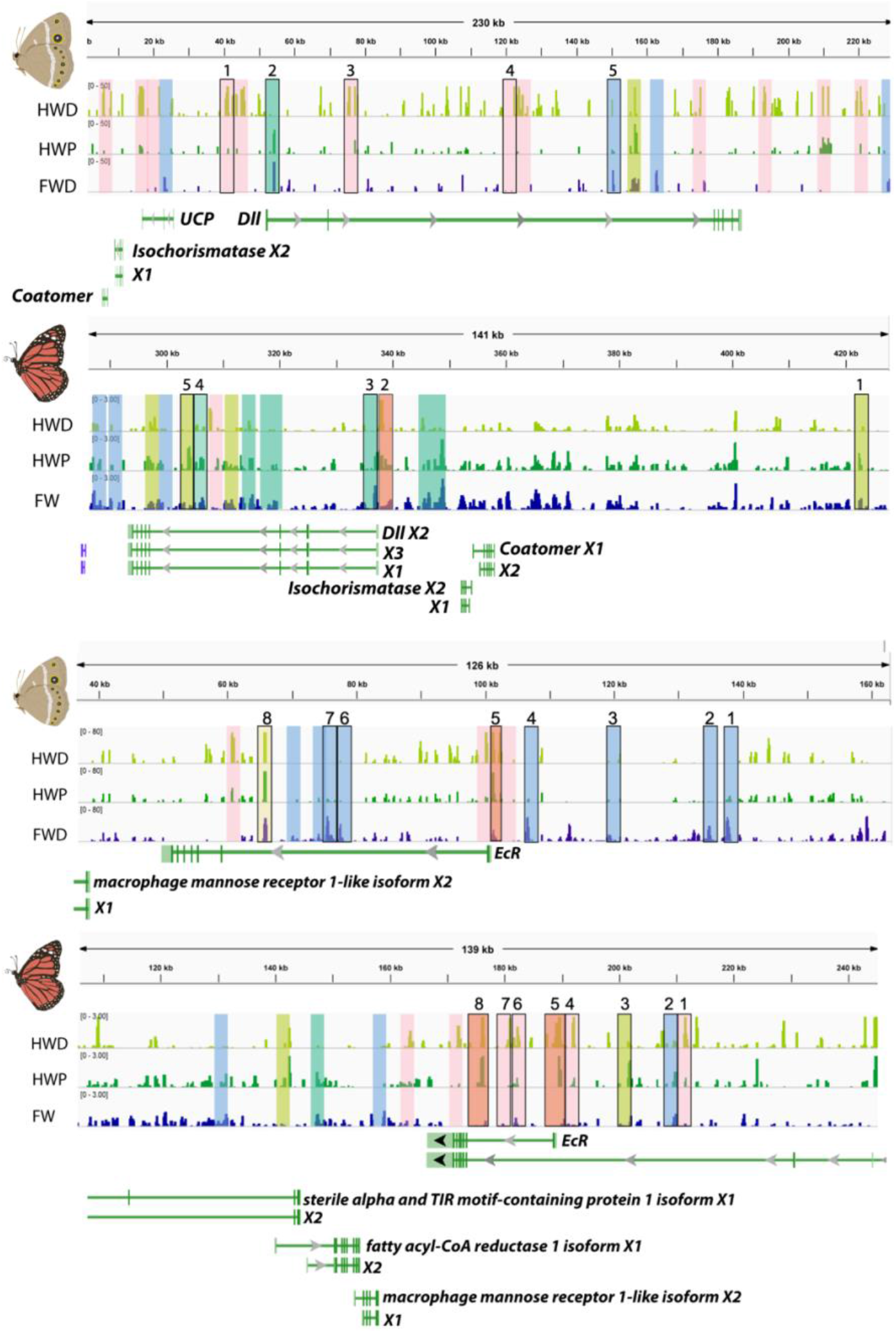
FAIRE-seq peaks for *B. anynana* and *D. plexippus* around *Dll* (top panel) and *EcR* (bottom panel). Significant peaks are colored as follows: Pink – HWD, Green – HWP, Blue – FWD/FW, Orange – overlaps between HWD and HWP, Mint green – overlaps between HWP and FWD/FW, Yellow – overlaps between all 3 wing regions, HWD+HWP+FWD/FW. Conserved peaks identified using mVista (minimum 50% conservation) are represented by shared numbers.

**Fig 3.**
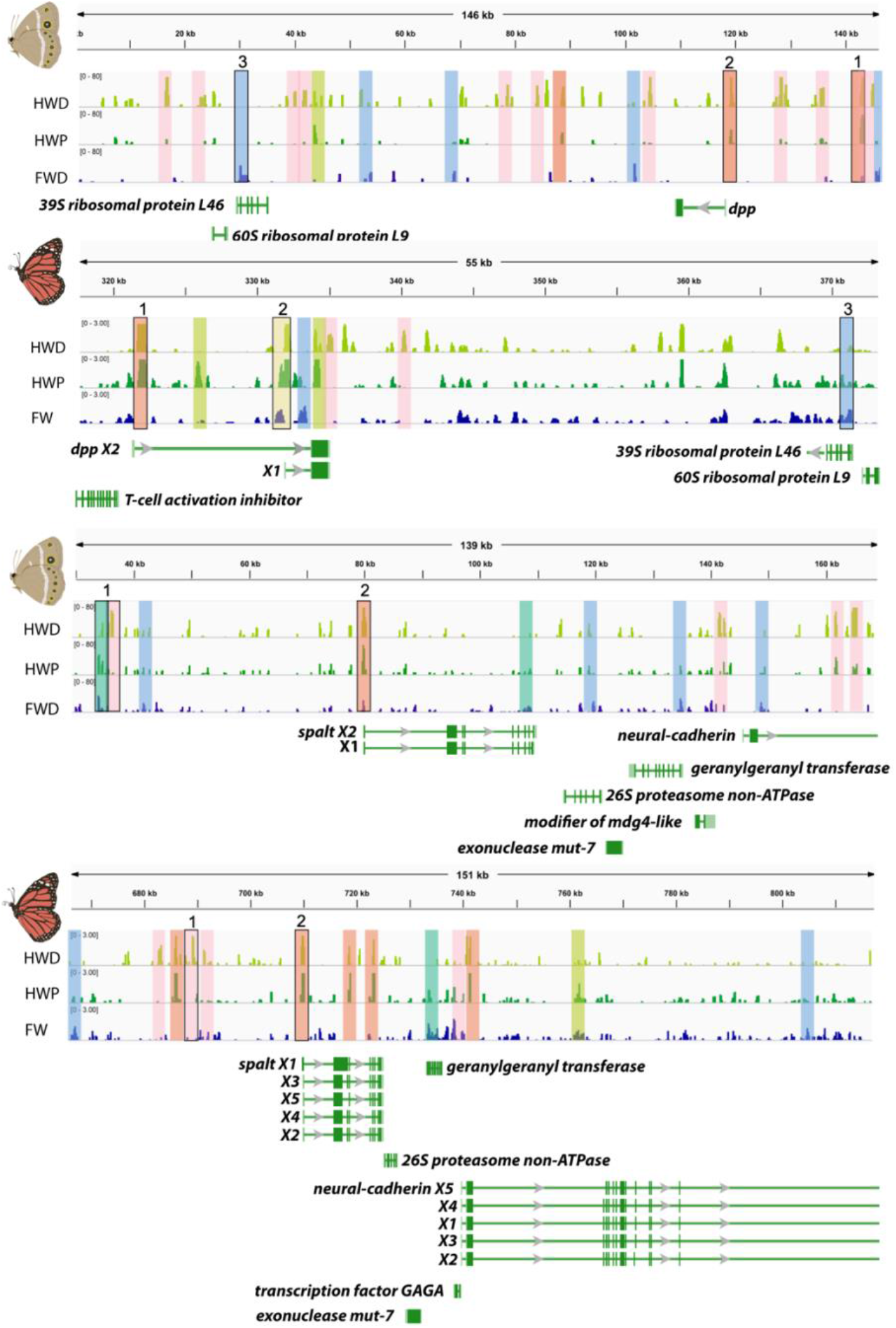
FAIRE-seq peaks for *B. anynana* and *D. plexippus* around *dpp* (top panel) and *spalt* (bottom panel). Significant peaks are colored as follows: Pink – HWD, Green – HWP, Blue – FWD/FW, Orange – overlaps between HWD and HWP, Mint green – overlaps between HWP and FWD/FW, Yellow – overlaps between all 3 wing regions, HWD+HWP+FWD/FW. Conserved peaks identified using mVista (minimum 50% conservation) are represented by shared numbers.

**Fig 4.**
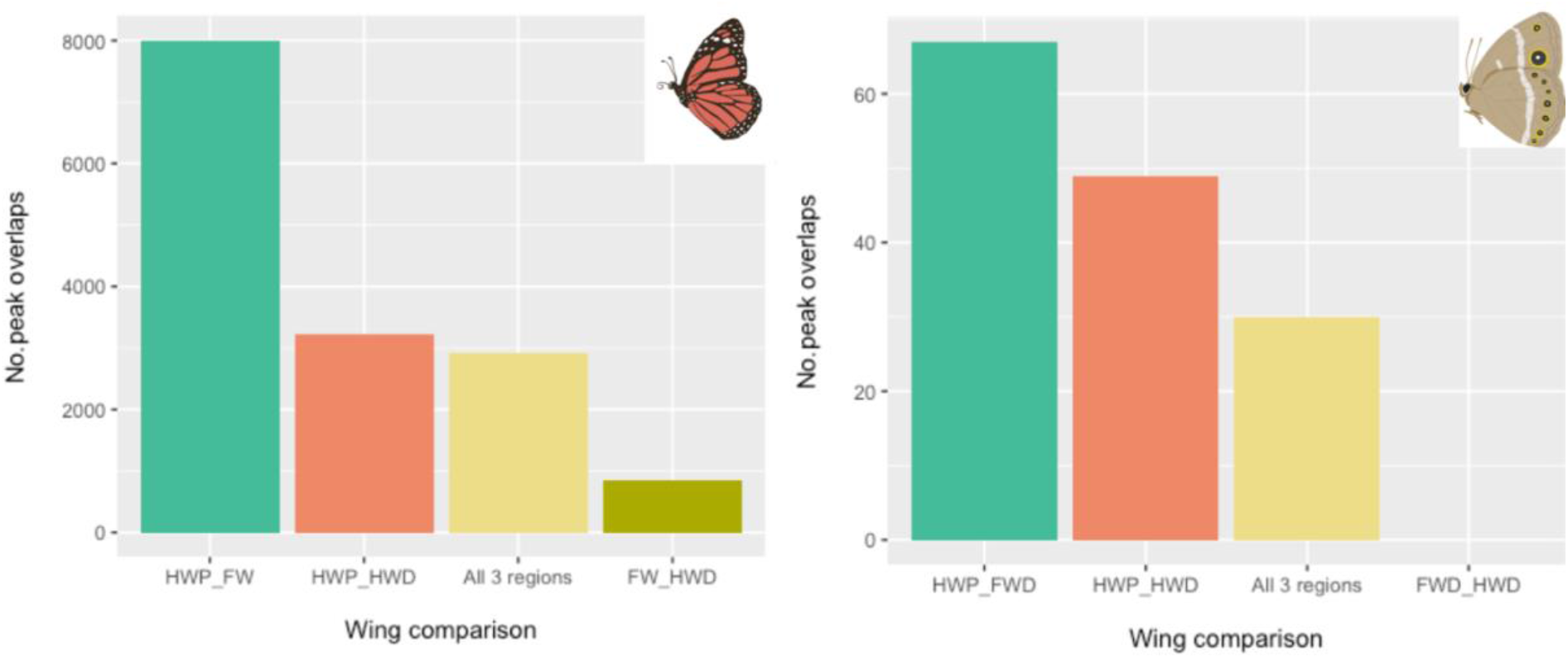
Number of overlapping OCRs between different wing regions for *D. plexippus* (based on OCRs extracted from the whole genome) and *B. anynana* (based on OCRs extracted from 31 scaffolds).

### Comparison of the chromatin landscape between *B. anynana* and *D. plexippus*

The two butterfly species have very different wing patterns and therefore we would expect to observe a different chromatin landscape even around the same genes. However, it is possible that some OCRs may be specific to particular wing regions and perhaps functionally conserved between both butterfly species. We first compared the chromatin profile around six genes known to be involved in wing patterning. In *B. anynana*, we observed 19 regions of open chromatin across the 231kb scaffold containing the *Distal-less* gene. Twelve of these peaks were found in the HWD only, four regions were open only in the FWD, one was open only in the HWP, and one open region was shared between the FWD and HWP region at the TSS (Fig 2). In contrast, in the same region around the *Distal-less* gene in *D. plexippus*, there was a single region of open chromatin in the HWD, however at the TSS there was also an overlapping peak between the FW and HWP region and another peak overlapping the HWD and HWP regions. Thus, in both butterflies there is a shared peak in the proximal hindwing and forewing tissue at the TSS. Comparing the open chromatin profiles around the remaining 5 genes, the profiles are largely different, however for *EcR* (Fig 2), *spalt* and *dpp* (Fig 3) the TSS of these genes overlaps with open chromatin in the HWD and HWP regions for both butterfly species. Chromatin profiles for *wingless* and *engrailed* are shown in Fig. S1.

### Most conserved peaks are open in different wing regions in the two butterfly species

Using mVista for genome alignments, we compared the scaffolds between these two butterfly species to identify regions of sequence conservation with a minimum threshold of 50%. We then examined whether any of these regions of sequence conservation overlapped regions of open chromatin, to look for conserved peaks between the two butterflies. In the region around *Distal-less* we identified 5 peaks which shared sequence conservation >50% between the two butterflies. In all 5 cases the conserved peaks were open in different wing regions in each butterfly. Interestingly, peak 2 which overlaps the TSS in both FWD and HWP libraries is not conserved with the region that overlaps the TSS in the FW and HWP libraries in *D. plexippus* but rather it shares sequence conservation with a different a peak in the HWD and HWP libraries. However, for *EcR* and *spalt* (Figs 2 and 3), the peaks at the TSS (peaks 5 and 2 respectively) are conserved and open in the same wing regions in both butterflies. For *EcR* and *dpp* (Figs 2 and 3), peaks 2 and 3 respectively, are also open in forewing tissue in both butterflies.

### Motif enrichment in regions of open chromatin

We next examined whether there was any evidence of motif enrichment in regions of open chromatin between the different wing regions. For *D. plexippus* we first pooled the HWD and HWP data allowing us to compare the FW and the HW. MEME-chip identified 50 motifs which were enriched in the FW versus the HW in *D. plexippus* (E-value: 1.2e-1458) (Fig S2A). The top hit matched to known motifs for the transcription factors, Jigr1 (Jing Interacting Gene Regulatory) and Bab1 (Bric-a-Brac). Significant secondary motifs were also identified using SpaMo (E-value: 1.58e-11). MEME-chip identified 47 motifs that were significantly enriched in the HW versus the FW (E-value: 7.0e-2424), with the top hit showing similarity to the transcription factors Mad (Mothers against dpp) and Crol (Crooked legs) (Fig S2B). Significant secondary motifs were also identified using SpaMo (E-value: 1.12e-13) which resemble the primary motif. Comparing OCRs across the hindwing, between HWD versus the HWP, MEME-chip identified 55 GC-rich motifs enriched in the HWD region with the top hit matching to Escargot and Trithorax-like (GAGA factor, GAF) (E-value: 5.8e-1652) (Fig S2C). The CentriMo feature in MEME-chip identified this motif as having a central distribution within the OCRs. Significant secondary motifs were also identified using SpaMo (E-value: 2.53e-8) which were also GC-rich. For the comparison between the HWP versus the HWD regions, 37 motifs were enriched in the HWP (E-value: 3.3e-564), an AT-rich motif matching Bab1 was identified as the top hit (Fig S2D). Significant secondary motifs were also identified using SpaMo which are also AT-rich.

For *B. anynana*, 10 motifs were identified as enriched in FWD versus HWD wings (E-value: 1.3e-003), 6 motifs were significantly enriched in HWD versus FWD (E-value: 8.8e-009) and 3 motifs were significantly enriched in HWD versus HWP (E-value: 4.0e-007). For each of these comparisons the top hits had no similarity to any known motif. For HWP versus HWD, 4 motifs were significantly enriched (E-value: 5.1e-004) with the top hit matching to the transcription factors Mad, Enhancer of split mβ, helix-loop-helix and odd paired.

### GC content of motifs and OCRs varies between different wings/regions

Based on the MEME-chip analysis, we observed that motifs enriched between different wings and regions appeared to differ in GC content. To examine whether there was a statistical difference in GC content we calculated the percent GC content for each motif for each wing/region in *D. plexippus* where we had the most data. We found that HW motifs had a significantly higher GC content than FW motifs (log_e_(W_Mann-Whitney_) = 3.87, r=-0.96, p<0.0001) (Fig 5A). We next looked to see if the GC bias for the HW motifs was explained by differences in GC content in the OCRs themselves. We found that HW OCRs had a significantly higher GC content than the FW OCRs (log_e_(W_Mann-Whitney_) = 18.26, r=-0.61, p<0.0001) (Fig 5B). The overall GC content of the genome is 28%, and the GC content of the FW OCRs is slightly lower at 24%. Both are much lower than the HW OCRs at 39% which is more similar to the coding regions at 46%. To examine whether the GC content is consistent across the HW itself, we repeated these analyses for the HWD and HWP OCRs. The motifs for the HWD OCRs had a significantly higher GC content than for the HWP OCRs (log_e_(W_Mann-Whitney_) = 7.53, r=-0.87, p<0.0001) (Fig 5C) and likewise the GC content of the HWD OCRs was significantly higher than the HWP OCRs (log_e_(W_Mann-Whitney_) = 18.22, r=-0.6, p<0.0001) (Fig 5D). Together these results show that HWD OCRs are enriched in GC content, whereas the HWP and FW OCRs display a much lower GC content which is reflected in the AT-enriched motifs. We next examined whether the higher GC content in the HWD region could be due to these OCRs being longer. We found that the average length of the OCRs was significantly different between all groups, however the FW had the longest OCRs and the HWP region had the shortest OCRs (KW= 419.7, r=-0.01, p<0.0001) (Fig S3A).

**Fig 5.**
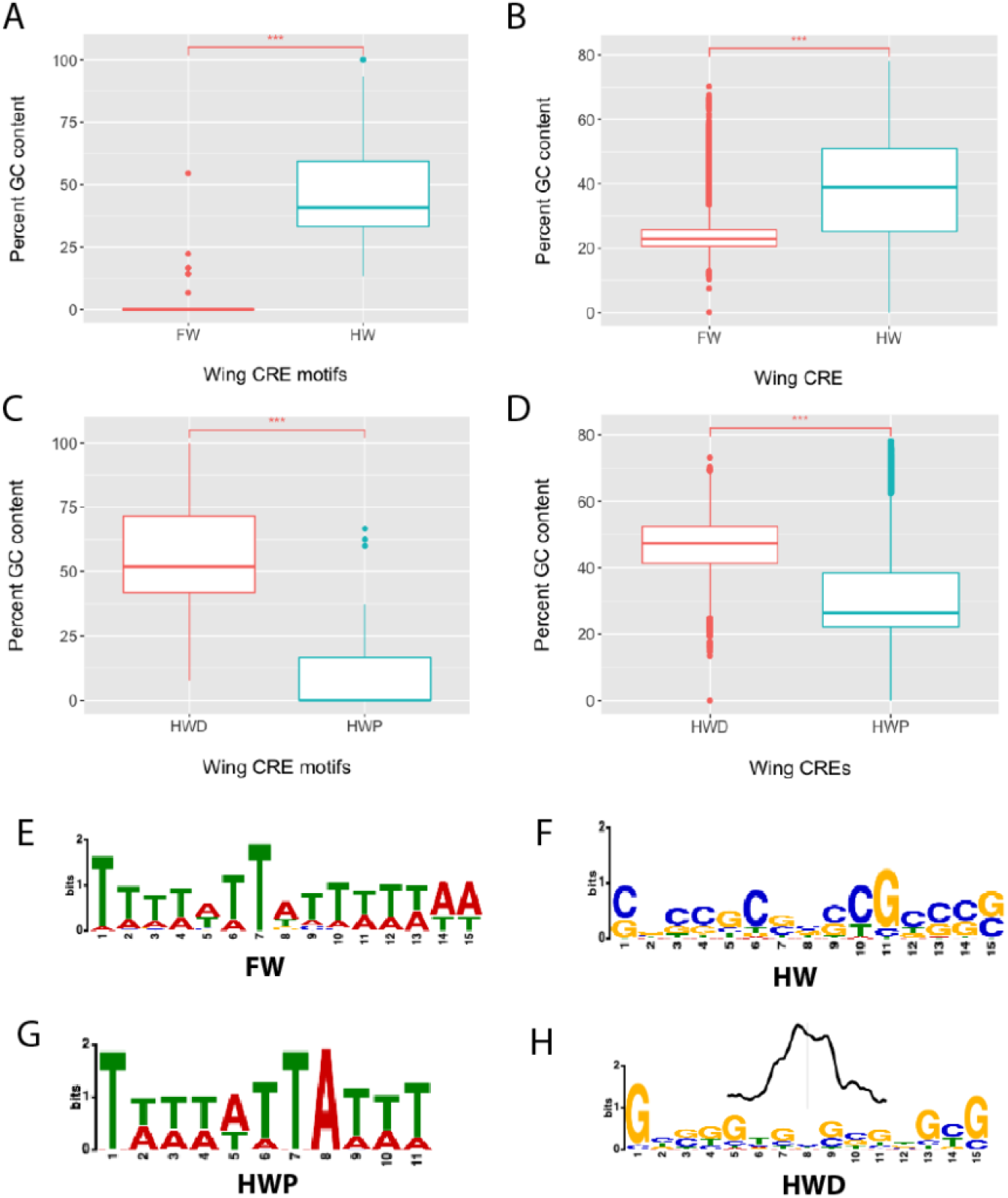
GC content analysis for *D. plexippus*. **A)** Boxplots showing average percent GC content in motifs in OCRs from forewings (Motifs: FW, n = 50) and hindwings (HW, n = 47) across the whole genome. **B)** Average percent GC content in OCRs in forewings (FW, n=21,111) and hindwings (HW, n=20,390) **C)** Average percent GC content. for motifs for HWD (n=55) and HWP (n=37) regions and **D)** in HWD (n=8,906) and HWP (n=11,484) OCRs across the whole genome. The box represents the interquartile range (IQR), the line represents the median, and the whiskers represent 1.5 times the IQR in the lower and upper quartiles. *** represents P<0.001, **E-H)** Top hit motifs for MEME using OCRs for the whole genome comparing **E)** FW versus HW – Jigr1/Bab1, **F)** HW versus FW – Mad/Crol, **G)** HWP versus HWD – Bab1, **H)** HWD versus HWP – Escargot/GAF with the centrimo graph generated by MEME-chip showing that this motif has a centralized position within the OCRs.

We next looked to see if this pattern held in *B. anynana* comparing the FWD with the HWD data. OCRs were extracted from all 31 scaffolds. Only a small number of motifs were found in the MEME-Chip analysis. The motifs enriched in the FWD OCRs had a GC content of 19% (n=13) compared to the motifs enriched in the HWD OCRs which had a much higher GC content of 69% (n=14). We also compared the GC content of the FWD and HWD OCRs and found that the GC content was significantly higher in the HWD (log_e_(W_Mann-Whitney_) = 9.47, r=-=0.46, p<0.0001) (Fig S4A). The combined scaffolds had a GC content of 32% compared to the whole genome which was 15% (5630 scaffolds). The GC content of the coding regions was much higher at 51%.

We also compared the OCRs between the HWD versus HWP regions. This analysis produced a small number of motifs, however in this comparison the HWP motifs (n=9) had a much higher GC content than the HWD motifs (n=6) at 86% versus 27% respectively. Comparing the GC content of the OCRs for these scaffolds, we observed a significantly higher GC content for the HWP than the HWD (t=-4.53, df=230.64, p<0.0001)(Fig S4B). The length of the OCRs were also significantly different for each wing region with the longest OCRs found in the HWD region and the shortest in the HWP region (KW= 30.71, r=-0.06, p<0.0001)(Fig S3B).

### Comparison of the GC content in open chromatin overlapping genomic elements

Finally, we examined whether the GC content varied in regions of open chromatin that overlapped different genomic elements to determine whether the differences in GC content were explained by OCRs open in specific parts of the genome (Fig S5-7). Focusing on the data from *D. plexippus*, we found that for each wing/region most of the open chromatin mapped to intergenic and intronic regions, and that the GC content showed significant differences between various genomic elements within each wing/wing region. While these differences were small, more variation was observed for the GC content in HWP OCRs that overlapped exons, the promoter and the TSS (Fig S6). Overall, we found that the GC content was, on average, similar across all genomic elements within each wing region, with a higher average in the distal hindwing.

## Discussion

We examined whether the chromatin landscape varies with wing pattern variation in two species of butterflies which exhibit some similarities and differences between forewings and hindwings and spatially across the hindwings. We hypothesized that in wing regions with similar color patterns such as the distal tissue with eyespots in *B. anynana*, the forewings and hindwings might share a similar chromatin profile. This hypothesis is based on previous work showing that the development of eyespots in forewings and hindwings is generally regulated by the same set of genes (Monteiro et al. 2015; Özsu et al. 2017; Connahs et al. 2019; Banerjee et al. 2020). Although the forewings and hindwings differ in the number of eyespots, we predicted that we might detect a common signature in the chromatin profile surrounding eyespot-associated genes at ~24 hours post-pupation. Our data however did not support this hypothesis. We did not identify any OCRs that were shared in both wing tissues with eyespots around the loci of 6 genes known to be important in eyespot development, nor were any overlaps found for the remaining 25 scaffolds. As we did not have FAIRE-seq data for the entire genome, we cannot rule out the possibility of shared OCRs between the distal forewing and hindwings in other regions of the genome.

Interestingly, although *D. plexippus* has a very different wing pattern with simple white spots along the distal margins of both wings, we found that that the fewest overlapping OCRs were observed between the distal hindwing and forewing tissue. As we profiled chromatin in the entire forewing we will have missed spatial differences in this wing, therefore the number of shared OCRs between the distal forewing and distal hindwing might be even smaller. Taken together our results do not show any evidence that similar phenotypes on different wing tissues share a similar chromatin profile at the same stage of development. Nor did our findings support those in *Junonia* and *Heliconius* where the chromatin profile was shown to be similar in forewings and hindwings at the same developmental stage regardless of phenotype (Lewis and Reed 2018a; Burg et al. 2019; Lewis, Geltman, Pollak, Rondem, and Belleghem 2019). Our findings suggest that there are distinct differences in how the chromatin is regulated between forewings and hindwings and that the distal wing regions, in particular, may exhibit low similarity.

Although we observed regional differences between the proximal and distal regions of the hindwing which differ morphologically for both butterflies, we also found OCRs that are shared across the entire hindwing including at the TSS. This could be due to the same genes being expressed across the entire wing tissue albeit at different levels in the proximal and distal regions. An unexpected finding was that most OCRs were shared between the distal forewing and proximal hindwing regions in *B. anynana* which show no morphological similarity. Similarly, we also found that the highest number of overlapping OCRs were between the forewing and proximal hindwing regions in *D. plexippus*. This observation is difficult to understand with our current data, and we later discuss some potential scenarios that may explain these patterns. Overall our results appear to be more in line with those from Bozek et al., (2019) who observed regional differences in open chromatin across the *Drosophila* embryo.

The discovery that many OCRs were unique to either forewings or hindwings may be explained by upstream regulators involved in differentiating the identity of these wings. The Hox gene *Ultrabithorax (Ubx)* is expressed only in hindwing tissue and knocking this gene out transforms hindwings to forewing identity in butterflies and moths (Matsuoka and Monteiro 2019; Tendolkar et al. 2021). Thus, *Ubx* could be involved in regulating the open chromatin profile that is unique to the hindwings. Loker et al., (2021) found that *Ubx* and its cofactors were important in regulating chromatin accessibility in *Drosophila* not only between the wing and haltere, but also spatially between the proximal and distal regions of the haltere. Their findings revealed a complex relationship between *Ubx* and its cofactors and how regional differences in the expression of these cofactors influence whether *Ubx* acts as a repressor or an activator of chromatin and gene transcription. Their findings also suggest that *Ubx* plays a key role in regulating chromatin dynamics in the T3 appendages which may explain why in *Heliconius* and *Junonia* the major signal in chromatin differences between forewings and hindwings was found at *Ubx* itself. *Ubx* and other Hox genes such as *Antennapedia* may have a similar role in butterflies, where spatial differences in expression of these genes and their collaborating transcription factors play an important role in regulating pattern development (Beh et al. 2016; Matsuoka and Monteiro 2019; Fang et al. 2021; Paul et al. 2021).

### Differential enrichment of transcription factor motifs across wings/regions

Given that Hox genes are important in specifying regional identity between forewings and hindwings (Deutsch 2005; Tomoyasu 2017; Paul et al. 2021) we looked for enrichment of motifs for Hox genes or other transcription factors that may be important in shaping morphological differences between the wings and wing regions. In *D. plexippus*, we found that a motif for the transcription factor Mad (Mothers against Dpp) and the zinc finger protein Crol (Crooked legs) was significantly enriched in the hindwing relative to the forewing. Surprisingly, despite the requirement of *Ubx* for hindwing identity we found no enrichment of Ubx-binding motifs. A ChiP-seq study against Ubx proteins in *Drosophila* haltere tissue also found no enrichment of Ubx motifs likely due to its low DNA binding specificity. Instead the study reported enrichment in motifs for GAF (GAGA binding factor) and Mad (Agrawal et al. 2011). The authors suggest that Hox proteins cooperate with other transcription factors in addition to their cofactors to achieve specificity. For example, Mad has previously been shown to collaborate with Ubx in the repression of *spalt* in halteres (Walsh and Carroll 2007). Whether Mad also collaborates with Ubx in butterfly wings is currently unknown.

Loker et al., (2021) also found no enrichment of the monomer Ubx-binding motif (TAAT)(jaspar.genereg.net/) when conducting a ChiP-seq analysis between the whole imaginal discs of the wing and haltere, however this enrichment was found in the haltere pouch when comparing the proximal and distal regions of the haltere, where Ubx binds alone in the haltere pouch. When we compared the distal and proximal hindwing regions, we found no enrichment of Ubx-binding motifs but instead found enrichment for a motif matching both Escargot and GAF-binding which was significantly enriched in the central region of the OCRs. Escargot regulates genes involved in wing development and can also function as a transcriptional repressor (Fuse et al. 1996). Whether Escargot interacts with Ubx however is unknown. GAF is known to regulate *Ubx* and functions as a pioneer factor involved in both activating and repressing transcription (Biggin and Tjian 1988; Chopra et al. 2008; Lomaev et al. 2017). When comparing the proximal with the distal hindwing region, Bab1 (Bric-a-brac) was significantly enriched in the HWP region. Bab1 interacts with a transcriptional regulator TAF3 (a component of the TFIID complex) which also binds to GAF – a potential cofactor of Ubx (Chopra et al., 2008; Agrawal et al., 2011). Future work could explore whether the different transcription factors enriched in each hindwing region are interacting with Ubx to differentially specify the wing color patterns.

In the forewing, motifs for the transcription factors, Jigr1 and Bab1 were significantly enriched relative to the hindwing. Not much is known about the function of Jigr1. In a study examining transcription factor cooperative binding, Kazemian et al., (2013) found that GAF and Jigr1 had the largest number of binding partners which could suggest that Jigr1 has a similar function to GAF in chromatin remodeling. Overall the motif analyses for all wings/wing regions suggests that the enriched motifs in the OCRs include binding sites for transcription factors potentially involved in chromatin remodeling.

### GC content of OCRs differs between wings and wing regions

One of the most striking observations from this analysis was that the different wings/regions were differentiated by the GC content of the transcription factor binding motifs. For both *D. plexippus* (FW versus HW) and *B. anynana* (FWD versus HWD), the forewings were characterized by AT-rich motifs whereas the hindwings were characterized by GC-rich motifs. We also found that the distal and proximal regions of the hindwing differed in the GC content of their motifs. For *D*.*plexippus*, the proximal hindwing region was characterized by AT-rich motifs in contrast to the distal hindwing region which was GC-rich, whereas the opposite trend was found for *B. anynana*. This difference in GC content of the motifs was explained by the GC content of the OCRs which appeared unrelated to their length. Distal hindwing OCRs had a higher GC content than the overall *D. plexippus* genome, however this was not due to their length as forewing OCRs were on average longer.

Several studies have shown that GC content is important in determining transcription factor binding. Transcription factors generally exhibit a specific preference not only for their binding motif but also for the sequence properties of the flanking regions (Dror et al. 2015). Some transcription factors prefer to bind at homotypic sites where the flanking regions have a similar GC content to the primary motif, however others prefer contrasting GC content in the 5’ and 3’ flanking regions (Dror et al. 2015; Dror et al. 2016; Yella et al. 2018). The GC content influences not only the propeller twist of the DNA, and its flexibility or rigidity but also the binding affinity of transcription factors which directly affects gene expression (Dror et al. 2016; Yella et al. 2018). Certain transcription families also show a preference for the GC content and structural features of the DNA. Zinc-finger transcription factors such as GAF, Crol, and Escargot show a preference for binding GC-rich regions whereas homeodomain containing transcription factors prefer AT-rich regions with a low propeller twist (Dror et al., 2016; Yella et al., 2018).

These differences in GC content in open chromatin between the wings and wing regions may represent a form of regulatory control determining which transcription factor complexes are able to bind. In the hindwing for example, these differences could limit the set of transcription factors that interact with Ubx to specify regional identity. As shown by Loker et al., (2021), the cofactors interacting with Ubx varied spatially across the haltere disc activating expression in certain regions yet repressing it in others. Very few ChiP-seq studies have been conducted in Lepidoptera (Burg et al., 2019; Lewis et al., 2016; Lewis et al., 2019b) thus more studies are required to identify spatial patterns of transcription factors binding to open chromatin in wing tissue.

### Sequence conservation of OCRs does not correspond to functional conservation

Although *D. plexippus* and *B. anynana* have very different wing patterns, we looked at the chromatin profile around six patterning genes to see whether OCRs that show >50% sequence similarity were open in the same wing or wing regions which could imply functional conservation. Overall, we found that many of the conserved OCRs were open in different wings/regions in each species. However, for *spalt, dpp* and *EcR*, the chromatin is open at the TSS across the entire hindwing in both species. Interestingly, Lewis et al., (2016) found that OCRs at the TSS were more conserved than distal OCRs across lepidopterans. Open chromatin at the TSS could represent expression of these genes. In *B. anynana*, Spalt proteins and *dpp* mRNA transcripts are present in the marginal eyespots during this developmental period (~24 hrs pp) (Banerjee et al. 2021), however we do not have any expression data for the pupal stages of *EcR* in *B. anynana* or for any genes in *D. plexippus*.

It is also difficult to know precisely which genes many of these OCRs regulate. We have previously used CRISPR to disrupt one of these conserved OCRs in an intron of *Dll* in *B. anynana* which led to loss of eyespots and disruption of other traits (Murugesan et al. 2022) mirroring our earlier work knocking out the *Dll* gene in this butterfly (Connahs et al., 2019). This OCR was open in the distal forewing in *B. anynana* yet it was open in the proximal hindwing in *D. plexippus*. Future work characterizing conserved OCRs open either in the same or different wing regions could provide insight into the evolution of their function. As our data represents a single time point, we also do not know whether these conserved OCRs share a similar temporal pattern of accessibility. Lewis et al., (2016) found that OCRs active across multiple developmental stages in butterflies showed increased sequence conservation. Thus, a temporal analysis or functional disruption would help to identify those conserved OCRs that are associated with a specific wing/wing region in both butterfly species which may suggest functional conservation. Taken together our results support previous studies suggesting that sequence conservation does not always imply functional conservation (Nelson and Wardle 2013).

### Limitations of the study

The main limitation of this study is that we only had one replicate for each library and thus it is possible that the variation we observed between the different wing regions was due to spurious results. However, we think this is unlikely as we observed a similar trend in both butterflies. In both species, we observed variation in the chromatin profile between the different wing regions, and we also observed that OCRs in forewing tissue were AT rich whereas the OCRs in hindwings were GC rich, suggesting that these observations may be a common feature in butterfly wings, however further work is required to validate these findings. Finally, we have confirmed that one of the OCRs identified from this FAIRE-seq data is indeed functional and that knocking it out using CRISPR, leads to phenotypes that would be expected from the disruption of a cis regulatory element that regulates *Distal-less* (Murugesan et al. 2022).

## Conclusions

Here, we have shown that the open chromatin profile varies between different wings and wing regions in two butterfly species revealing how the chromatin landscape can change spatially even within the same tissue. We did not however find a clear association between the chromatin profile and the wing color patterns as demonstrated with the shared OCRs between the distal forewing and proximal hindwing of *B. anynana*. Our results and those of others highlight the importance of understanding not only the chromatin profile but also the transcription factors that are bound to those OCRs. Although the same OCRs may be open in different wing regions, the binding of transcription factors functioning as activators of nearby genes in one context and as a repressors in another, even in the same tissue, may explain spatial variation in color patterns. Further work is required to understand the role of Hox genes and their cofactors, as well as of other pioneer transcription factors, in generating OCRs, and how structural features of DNA may influence the binding preferences of these and other factors to regulate morphological variation.

## Supporting information

Bicyclus anynana MACS2 data

Danaus plexippus MACS2 data

## Acknowledgements

We thank Christine Merlin, at Texas A&M University, for sending us Monarch eggs. This work was supported by National Research Foundation (NRF), Singapore under its Investigatorship programme (award NRF-NRFI05-2019-0006).

## Author contributions

AM and MDG designed the experiment, MDG prepared the samples for FAIRE-seq, HC analyzed the data and wrote the manuscript. AM edited the manuscript.

## Methods

Wild-type *B. anynana* were maintained in lab populations and reared at 27°C and 60% humidity inside a climate room with 12:12 h light:dark cycle. *D. plexippus* eggs were obtained from Christine Merlin, at Texas A&M University, and were reared in the same climate chambers. *Bicyclus* larvae were supplied with young corn leaves and *Danaus* with tropical milkweed plants until pupation.

Wings from both species were dissected at ~22-26 hours post-pupation. FAIRE-enriched libraries for *D. plexippus* included 3 libraries, one prepared from whole forewings, and two from partial hindwings (proximal and distal regions). For the control input library (non-enriched), two whole forewings and two whole hindwings were pooled (Fig.1). For *B. anynana*, three FAIRE-enriched libraries were prepared including a forewing distal library which had its own control using distal forewing tissue only, and two from partial hindwings (proximal and distal regions) with control inputs of two whole hindwing tissues. All FAIRE-enriched libraries were prepared from 7-8 pooled wing tissues. For this experiment we only collected one replicate per library.

Libraries were prepared by Genotypic Technology (India), as paired-end reads (75*2) and sequenced using Illumina NextSeq. Quality checking of the raw reads was performed using FASTQC v0.11.3 and reads which had a phred score>30 were retained for downstream analyses. The reads were aligned to reference scaffolds or genomes for each species (BACs for *B. anynana* and the whole genome for *D. plexippus*) with BWA (0.7.13) using the following parameters –k INT, -w INT, -A INT, -B INT, -O INT, -E INT, -L INT, -U INT. The SAM files were converted to BAM files using SAMtools-0.1.7a and the resulting BAM files were converted to sorted BAM followed by removal of PCR duplicates. The final BAM files were then converted to BEDgraph files using BEDtools-2.14.3. Peaks were called with the MACS2 software using the aligned enriched and input (control) files with the qvalue (minimum FDR) cutoff to call significant peaks. Fold-enrichment and log likelihood scores were calculated using the command bdgcmp script on the enriched and input BEDgraph files. The bdgcmp command also removes noise from the enriched sample relative to the control. The BEDgraph files were converted BigWig files using bdg2bw for visualization in Integrative Genome Viewer (IGV) to identify the peaks (Thorvaldsdóttir et al. 2013). The BigWig files underlying this article are available in the Dryad Digital Repository which can be found at doi:10.5061/dryad.rv15dv492. The data showing all the peak calls from MACS2 can be found in the supplementary excel files S1-6.

## Identifying conserved non-coding regions

To identify conserved non-coding regions between *B. anynana* and *D. plexippus*, we used published BAC (Bacterial Artificial Chromosome) sequences from *B. anynana* around six genes known to be involved in eyespot pattern development (*Distal-less, EcR, spalt, dpp, wingless* and *engrailed*), the patterns found in the distal portions of wings (Conceição et al., 2011). The *B. anynana* BACs were approximately 200kb so to extract the same regions in *D. plexippus* we used Ensemble (ensembl.org) to identify the correct genomic location and export 100kb up and downstream of each gene. The genome region was exported along with the annotation files in vista format for use in mVista (Frazer et al. 2004). mVista was used to identify conserved regions between *B. anynana* and *D. plexippus* using the Shuffle-Lagan algorithm, which can identify rearrangements, duplications and transposition events and improve alignment of distant homologues (Brudno et al. 2003). Softmasking was used on *B. anynana* sequences and sequences were reverse-complemented where appropriate. Conservation parameters were changed to a minimum of 50% identity. The conserved sequences obtained from mVista along with the FAIRE-seq regions were annotated in geneious version 9.1 (Kearse et al. 2012) to identify overlaps between the conserved regions and the FAIRE-seq peaks.

## Motif enrichment and GC content analysis

MEME-chip was used to examine motif enrichment in the regions of open chromatin (Ma et al. 2014). MEME-chip is typically used to examine motif enrichment in 100bp peak regions in chip-seq data, however it can also be used locally for analysis of open chromatin by modifying the - ccut parameter from 100 to 0. This ensures that the whole region of open chromatin is examined rather than just the central 100 bp. Regions of open chromatin were extracted from each butterfly genome using bedtools. The databases used for MEME-chip were dmmpmm2009.meme, fly_factor_survey.meme, flyreg.v2.meme and /idmmpmm2009.meme (see S8 for full command line used). The output from MEME-chip produces a motif alignment file which was used to extract the motifs for GC content analysis. Prior to the analysis, duplicate motifs were removed, and the motif file was converted to a fasta file. GC content was examined using a python script (see S8 for the command line used). To examine shared regions of open chromatin between different wings and wing regions, bed files were created for each wing group comparison and the command line program Bedops (Neph et al. 2012) was used to generate peak overlap data with a minimum overlap set at 50 bp (see S8 for the command line used). All statistical analyses were performed in Rstudio (RStudio 2020).

