## Supplementary materials for "Butterfly wings exhibit spatial variation in chromatin accessibility"

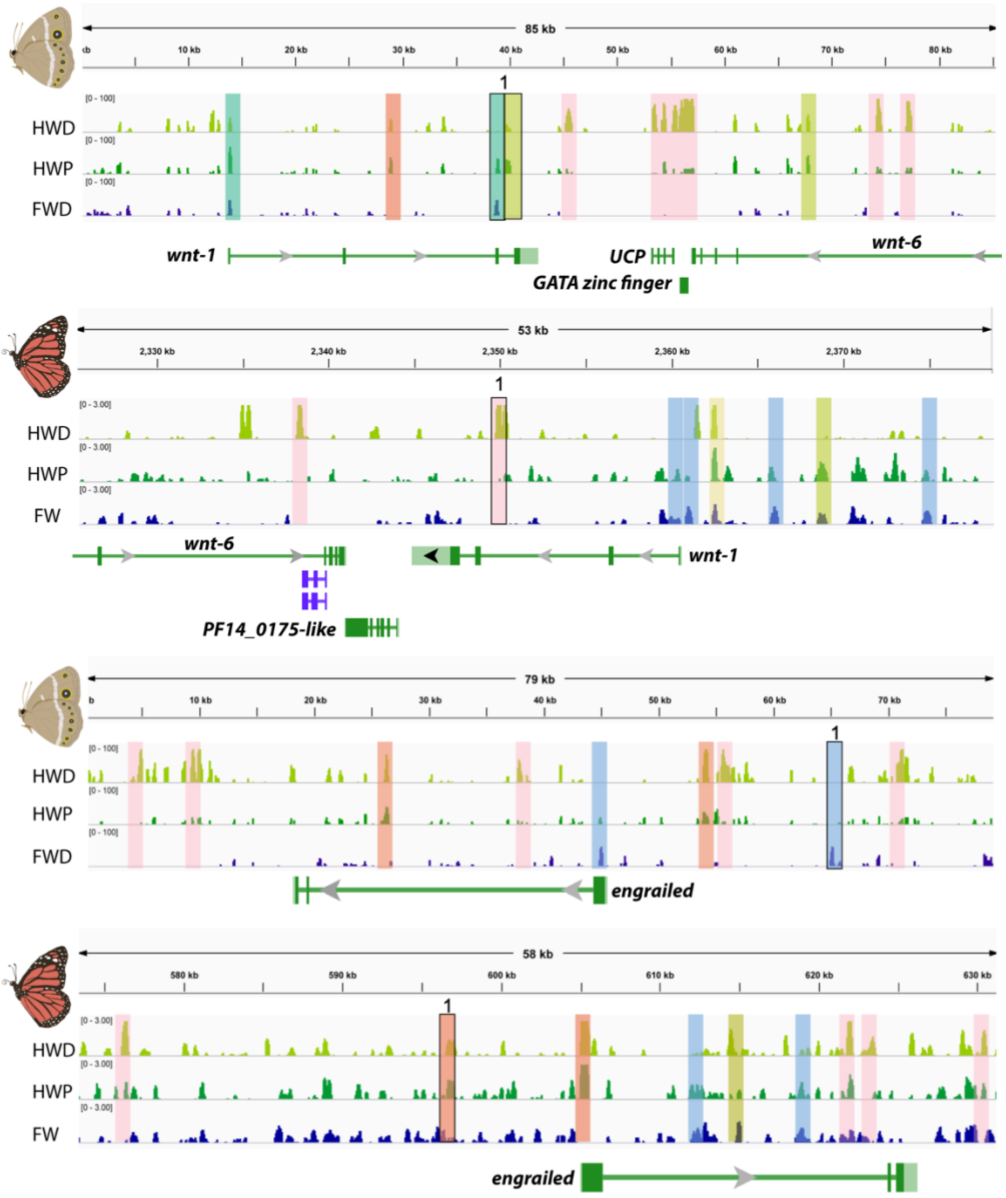


**Fig S1. Faire-seq peaks for *B. anynana* and *D. plexippus* around *wg* (top panel) and *engrailed* (bottom panel).** Significant peaks are colored as follows: Pink – HWD, Olive green, overlaps between HWD and FW. Green – HWP, Blue – FWD/FW, Orange – overlaps between HWD and HWP, Mint green – overlaps between HWP and FWD/FW, Yellow – overlaps between all 3 wing regions, HWD+HWP+FWD/FW. Conserved peaks identified using mVista (50%+ conservation) are represented by shared numbers.

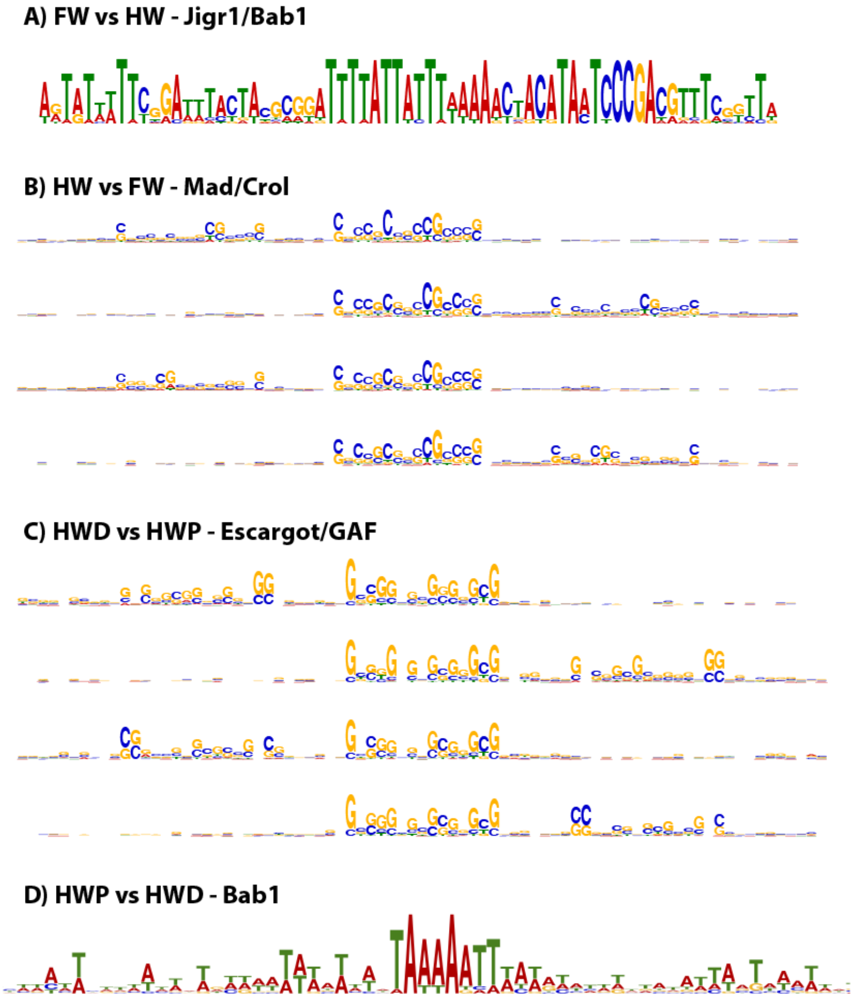


**Fig S2.** MEME-Chip results for enriched transcription factor binding-motifs in each wing/region with SpaMo results showing the flanking secondary motifs in OCRs from *D. plexippus*

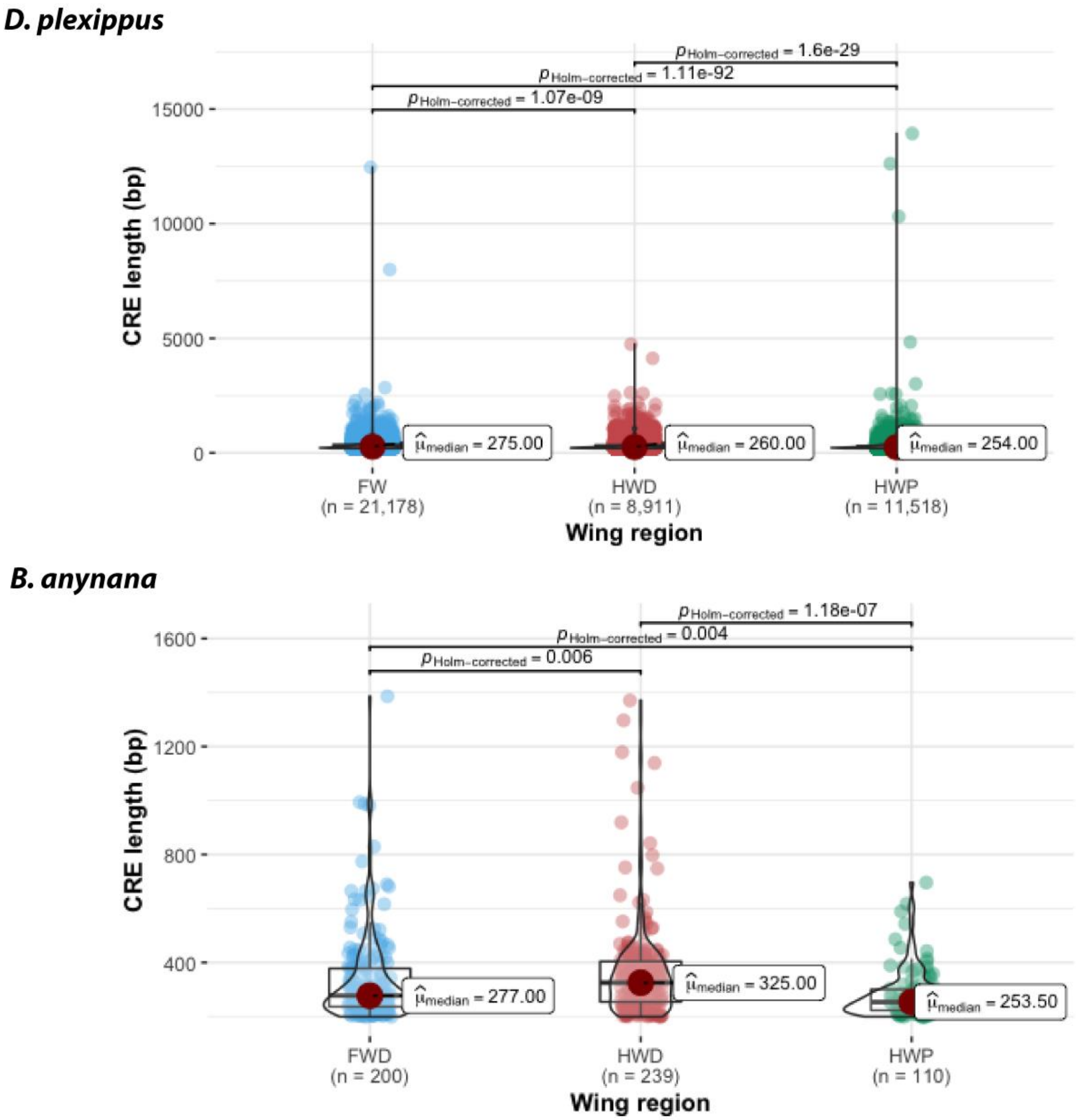


**Fig. S3.** Violin-boxplots showing the average OCR length for *D. plexippus* and *B. anynana* for the different wing regions. The box represents the interquartile range (IQR), line represents the median, and the whiskers represent 1.5 times the IQR in the lower and upper quartiles.

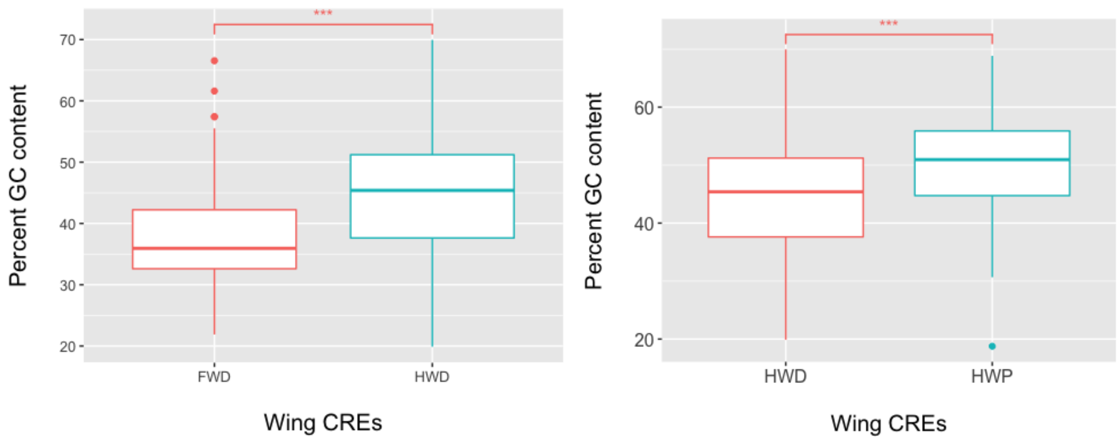


**Fig S4. A)** Boxplots showing the average percent GC content for *Bicyclus anynana* forewing and hindwing distal OCRs across 31 scaffolds, FWD: n=200, HWD: n = 239. **B)** Average percent GC content for *Bicyclus anynana* hindwing distal and hindwing proximal OCRs across 31 scaffolds, HWD: n=239, HWD: n = 110.The box represents the interquartile range (IQR), line represents the median, and the whiskers represent 1.5 times the IQR in the lower and upper quartiles. *** represents P<0.001

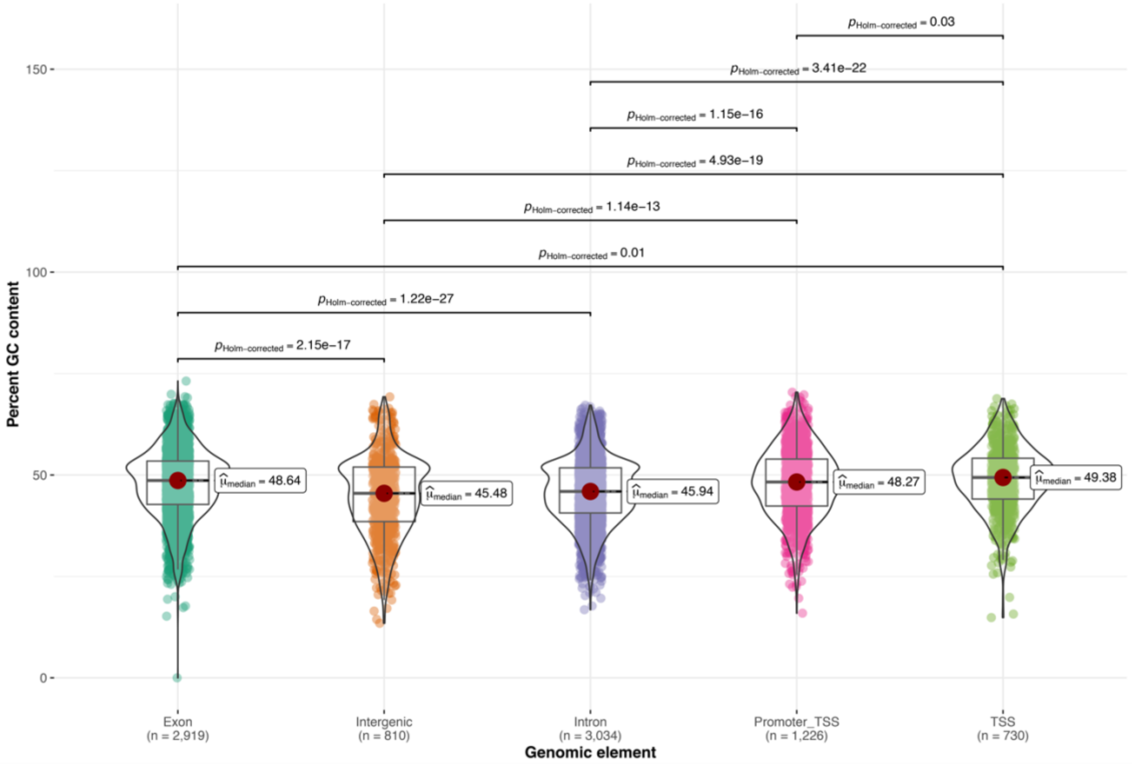


**Fig S5.** Violin-Boxplots for the distal hindwing (HWD) showing the average percent GC content in open chromatin overlapping different genomic elements for each wing region for *D. plexippus*. The box represents the interquartile range (IQR), line represents the median, and the whiskers represent 1.5 times the IQR in the lower and upper quartiles. Only the comparisons which are significantly different are annotated.

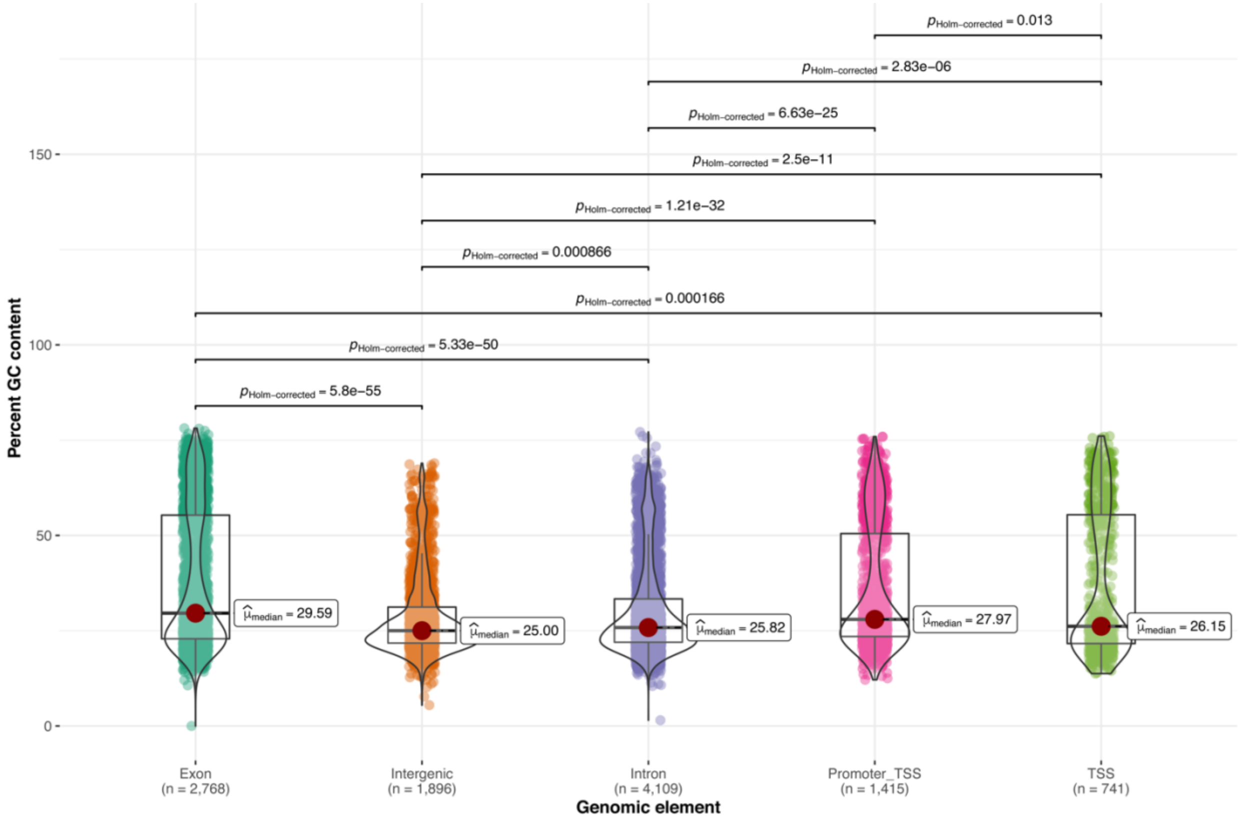


**Fig S6.** Violin-Boxplots for the proximal hindwing (HWP) showing the average percent GC content in open chromatin overlapping different genomic elements for each wing region for *D. plexippus*. The box represents the interquartile range (IQR), line represents the median, and the whiskers represent 1.5 times the IQR in the lower and upper quartiles. Only the comparisons which are significantly different are annotated.

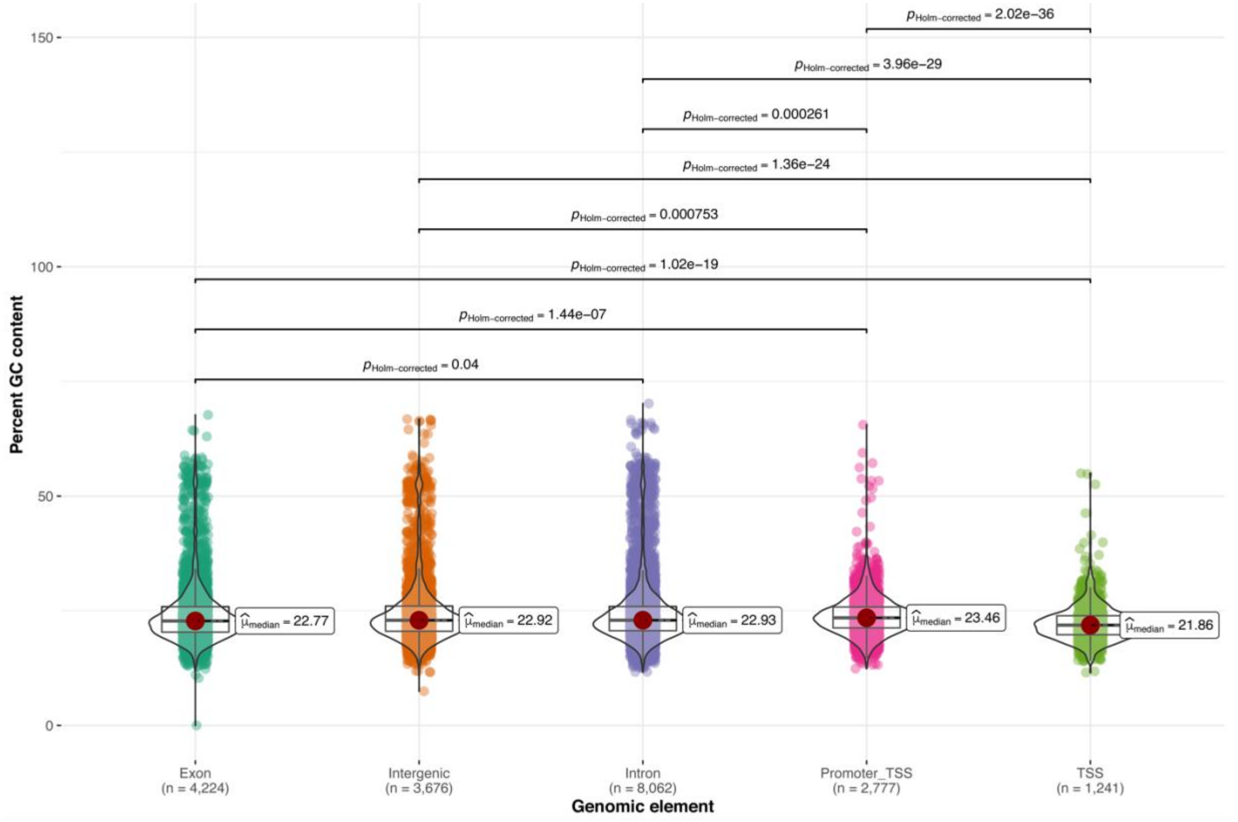


**Fig S7.** Violin-Boxplots for the forewing (FW) showing the average percent GC content in open chromatin overlapping different genomic elements for each wing region for *D. plexippus*. The box represents the interquartile range (IQR), line represents the median, and the whiskers represent 1.5 times the IQR in the lower and upper quartiles. Only the comparisons which are significantly different are annotated.

**S8 Command line scripts used:**

**Sample command line for MEME-chip**

```
meme-chip -oc ./Genome_OCRs_FW_vs_HW -time 240 -ccut 0 -dna -neg
Monarch_All_OCRs_HW.fasta -order 2 -minw 6 -maxw 15 -db
db/FLY/OnTheFly_2014_Drosophila.meme -db db/FLY/dmmpmm2009.meme -db
db/FLY/fly_factor_survey.meme -db db/FLY/flyreg.v2.meme -db
db/FLY/idmmpmm2009.meme -meme-mod zoops -meme-nmotifs 3 -meme-searchsize 100000 -
streme-pvt 0.05 -streme-totallength 4000000 -centrimo-score 5.0 -centrimo-ethresh 10.0
Monarch_All_OCRs_FW.fasta
```

**Commands for peak overlaps**

```
bedops --everything sorted_Monarch_HWD_overlaps.bed
sorted_Monarch_FW_overlaps.bed >sorted_MasterList_HWD_FW.bed

bedmap --echo --count --bp-ovr 50 --echo-map-id sorted_MasterList_HWD_FW.bed
sorted_MasterList_HWD_FW.bed  > Masteroverlaps_HWD_FW.tsv
```

**Python script for calculating GC content**

**https://warwick.ac.uk/fac/sci/moac/people/students/peter_cock/python/fasta_n/**

```
input_file = open(file.fasta', 'r')
output_file = open(file.tsv','w')
output_file.write('Gene\tA\tC\tG\tT\tLength\tCG%\n')
from Bio import SeqIO
for cur_record in SeqIO.parse(input_file, "fasta") :
#count nucleotides in this record...
	gene_name = cur_record.name
	A_count = cur_record.seq.count('A')
	C_count = cur_record.seq.count('C')
	G_count = cur_record.seq.count('G')
	T_count = cur_record.seq.count('T')
	length = len(cur_record.seq)
	cg_percentage = float(C_count + G_count) / length
	output_line = '%s\t%i\t%i\t%i\t%i\t%i\t%f\n' % \
	(gene_name, A_count, C_count, G_count, T_count, length, cg_percentage)
```

```
        output_file.write(output_line)
output_file.close()
input_file.close()
```
